# Synaptic Plasticity as a Function of the Temporal Derivative

**DOI:** 10.64898/2026.06.05.730489

**Authors:** Jinyoung Jang, Juan C. Flores, Karen Zito, Randall C. O’Reilly

## Abstract

A major outstanding question in neuroscience is whether the neocortex uses the same powerful learning algorithm as current AI models: error backpropagation. One way this could be accomplished is as a function of the temporal derivative (i.e., differences in neural activity states over time), which can closely approximate the backpropagated error gradient. We tested the hypothesis that the direction of synaptic plasticity is a function of the temporal derivative in synaptic activity over the course of a 200 ms (5 Hz) theta cycle. Using mouse hippocampal slices, we drove presynaptic activity across the two 100 ms halves of a 200 ms window at either 25 Hz or 50 Hz, combined with corresponding low and high magnitudes of postsynaptic depolarization, testing all four 2×2 combinations of these low and high activity levels, while measuring the resulting effects on synaptic efficacy (as measured by EPSP amplitude to standard test probes). Consistent with the computational hypothesis, a positive temporal derivative (low to high) resulted in LTP (increased synaptic strength), while a negative temporal derivative (high to low) resulted in LTD. Critically, both no-change conditions (stable low or high across 200 ms) resulted in no net synaptic change, even though the high no-change condition had the highest overall synaptic activity levels. Possible biochemical mechanisms that could support these results are discussed.

## Introduction

Understanding how the neocortex learns is perhaps the single most important step in understanding human intelligence, because our cognitive functions emerge over years of experience-driven learning within this brain structure, which is unique to mammals and is most greatly expanded in primates, especially humans. Current artificial intelligence (AI) systems are based on the powerful error-backpropagation learning algorithm, which has long been recognized as the single most capable learning mechanism in artificial neural networks (Rumelhart et al., 1986; Widrow & Hoff, 1960; Werbos, 1974). Thus, this algorithm provides the best computational-level hypothesis for how the neocortex should learn.

There have been a variety of proposals for how error backpropagation could be implemented in the brain (Lillicrap et al., 2020). Here, we test the predictions of one of the earliest such proposals, which is based on the idea that the backpropagated error gradient can be approximated by the *temporal derivative* in neural activity states over time (Ackley et al., 1985; Movellan & McClelland, 1993; O’Reilly, 1996; Xie & Seung, 2003; Scellier & Bengio, 2017). Specifically, in a network with bidirectional connectivity, which enables activation to flow in both bottom-up and top-down directions, changes in neural activity across any subset of neurons within the network will reverberate throughout the rest of the network. If these changes represent the difference between a prediction versus the correct outcome, and they can drive local synaptic plasticity according to the temporal derivative, then the resulting learning approximates error backpropagation.

Figure 1 illustrates this in the context of a simple three-layer network, with two distinct *phases* of neural activity, starting with an initial *prediction* or *minus* phase that reflects the impact of a given *input* pattern presented over the Sensory Input layer of simulated neuron-like processing units. Subsequently, the *outcome* or *plus* phase of activity arises when the *actual outcome* (i.e., correct or target) activity pattern is driven onto the Prediction layer. An error signal can be computed via the simple subtraction of these activity states: (*plus – minus*) or (*outcome – prediction*), i.e., the *temporal derivative* or *temporal difference*, at any neuron anywhere. This error signal provides a good approximation to the error gradient that would otherwise be computed by error backpropagation (O’Reilly, 1996).

**Figure 1.**
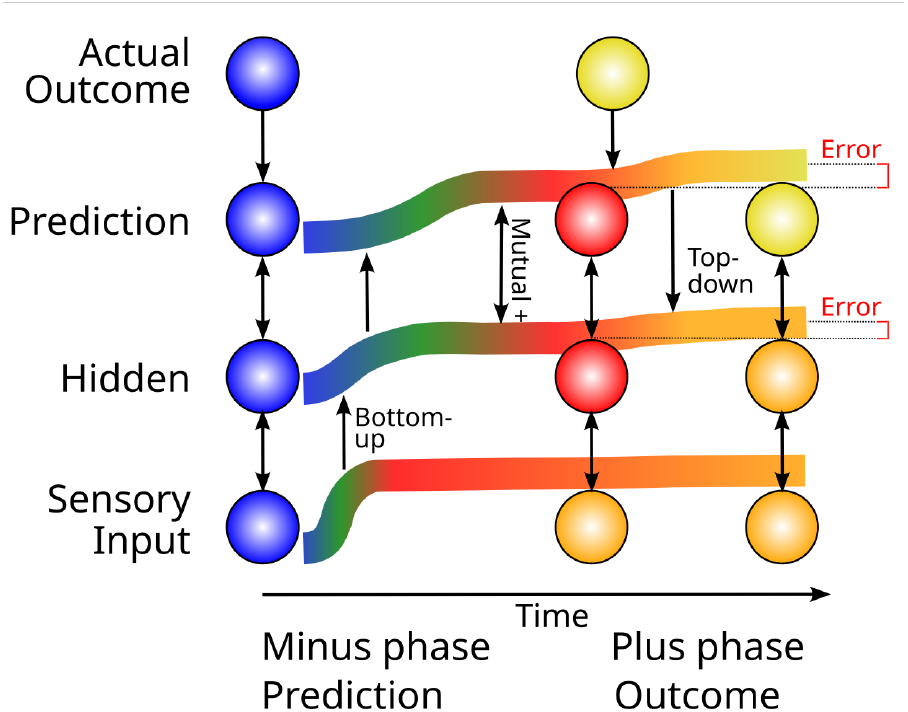
How bidirectional activation propagation can communicate error signals, in the simplest case of a three-layer network mapping from a Sensory Input to a Prediction output, with the Actual Outcome driving the Prediction layer only in the later *plus* phase, after an initial *minus* phase when the prediction is generated. The Error is the temporal difference between the (*plus* – *minus*) activity levels. There is just one network with three bidirectionally connected units, as shown at the left; the networks shown further to the right are snapshots of the activity state of this network at different points in time, which evolves from left to right. The thick colored lines also show the activation level of each of the three neurons over time, both in terms of the line height and the brightness and warmth of the color gradient. Initially, each neuron is inactive (blue). Then, external Sensory Input arrives, and a wave of bottom-up excitation propagates upward through the Hidden and Prediction layers. Critically, the Prediction and Hidden neurons mutually excite each other via bidirectional connections, which contributes to each of their activity levels. The snapshot of the network in the middle shows the neural activity at the end of the minus phase. Then, at the start of the plus phase, the Actual Outcome arrives, which is more active than the Prediction, and it therefore drives more activity in the Prediction neuron. This propagates top-down to the Hidden neuron as well, which is the key mechanism by which bidirectional connectivity communicates error signals, causing the Hidden neuron to have a (*plus* – *minus*) activity difference, reflecting the top-down influence from the Prediction layer. This temporal-difference based Error signal provides a good approximation to the error backpropagation error gradient (O’Reilly, 1996).

Thus, the direct biological prediction from this type of error-driven learning is that the direction of synaptic plasticity should be a function of the change in activity (i.e., temporal derivative) across a time window that would encompass this transition between the prediction and outcome phases. Note that despite both being based on changes over time, this neocortical learning mechanism is entirely distinct from the *TD* (*temporal difference*) reinforcement learning algorithm that describes the behavior of dopamine neurons in the midbrain (Sutton & Barto, 1998; Montague et al., 1996). In TD, dopamine neurons represent the temporal difference *explicitly* in their firing rates. By contrast, in neocortical temporal derivative learning the error gradient remains *implicit* in the changes in neural firing over time, and yet this temporal derivative drives synaptic plasticity locally everywhere. This implicit representation of the error gradient has critical advantages in simplifying neural computation as elaborated in the discussion.

A further elaboration of this learning algorithm provides a specific hypothesis regarding the duration of this time window, and a biologically explicit hypothesis regarding the source of the prediction and outcome signals driving this form of learning (O’Reilly et al., 2021). Specifically, unique features of the thalamocortical circuitry between the neocortex and the pulvinar nucleus of the thalamus should drive an alternating sequence of prediction-then-outcome states (Figure 2), over the course of a 200 ms (5 Hz) theta cycle. This hypothesis is consistent with considerable evidence at multiple levels, as reviewed in O’Reilly et al. (2021) (e.g., Fiebelkorn & Kastner, 2021; Sherman & Guillery, 2006; Sherman & Usrey, 2024). Furthermore, this same theta-cycle temporal derivative learning mechanism also applies to learning in area CA1 of the hippocampus (Ketz et al., 2013; Zheng et al., 2022).

**Figure 2.**
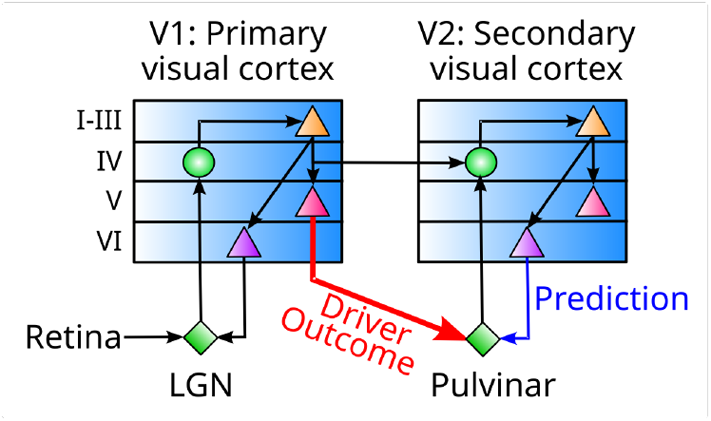
Connectivity between the neocortex and the pulvinar nucleus of the thalamus, in the case of primary and secondary visual areas, that is uniquely well-suited for driving predictive error-driven learning. The numerous and relatively weaker projections from layer 6 (VI) neurons activate a prediction over the pulvinar, that integrates the signals from multiple cortical areas and neurons to synthesize the prediction, which improves over the course of learning throughout the neocortex and in these final projections into the pulvinar. By contrast, the strong, focal driver inputs from layer 5 (V) intrinsic bursting (5IB) neurons can activate an outcome representation that is essentially an unlearned copy of the activity pattern in lower cortical layers (e.g., V1 trains V2 predictions in this case). The periodic bursting of the 5IB neurons ensures that this outcome activity is only phasically present (i.e., the plus phase), with a complete prediction – outcome learning cycle occurring within roughly 200 ms (i.e., theta frequency, 5 Hz). Diagram based on Sherman & Guillery (2006).

To test whether synaptic plasticity in the brain might be sensitive to the temporal derivative over a period of roughly 200 ms, we used a standard experimental preparation in mouse-brain slices that includes area CA1 and its afferent axonal fibers that originate in area CA3, which has been removed. Currents from individual CA1 neurons were recorded in whole-cell mode and postsynaptic neurons were stimulated using current injection, while the axonal afferents were also stimulated using bipolar electrodes, providing precise experimental control over the level of activity over time at the synaptic inputs to these CA1 neurons. We manipulated the level of activity in the synaptic inputs and the clamped postsynaptic CA1 neuron in a coordinated manner, across the two 100 ms halves of the 200 ms theta cycle (Figure 3).

**Figure 3.**
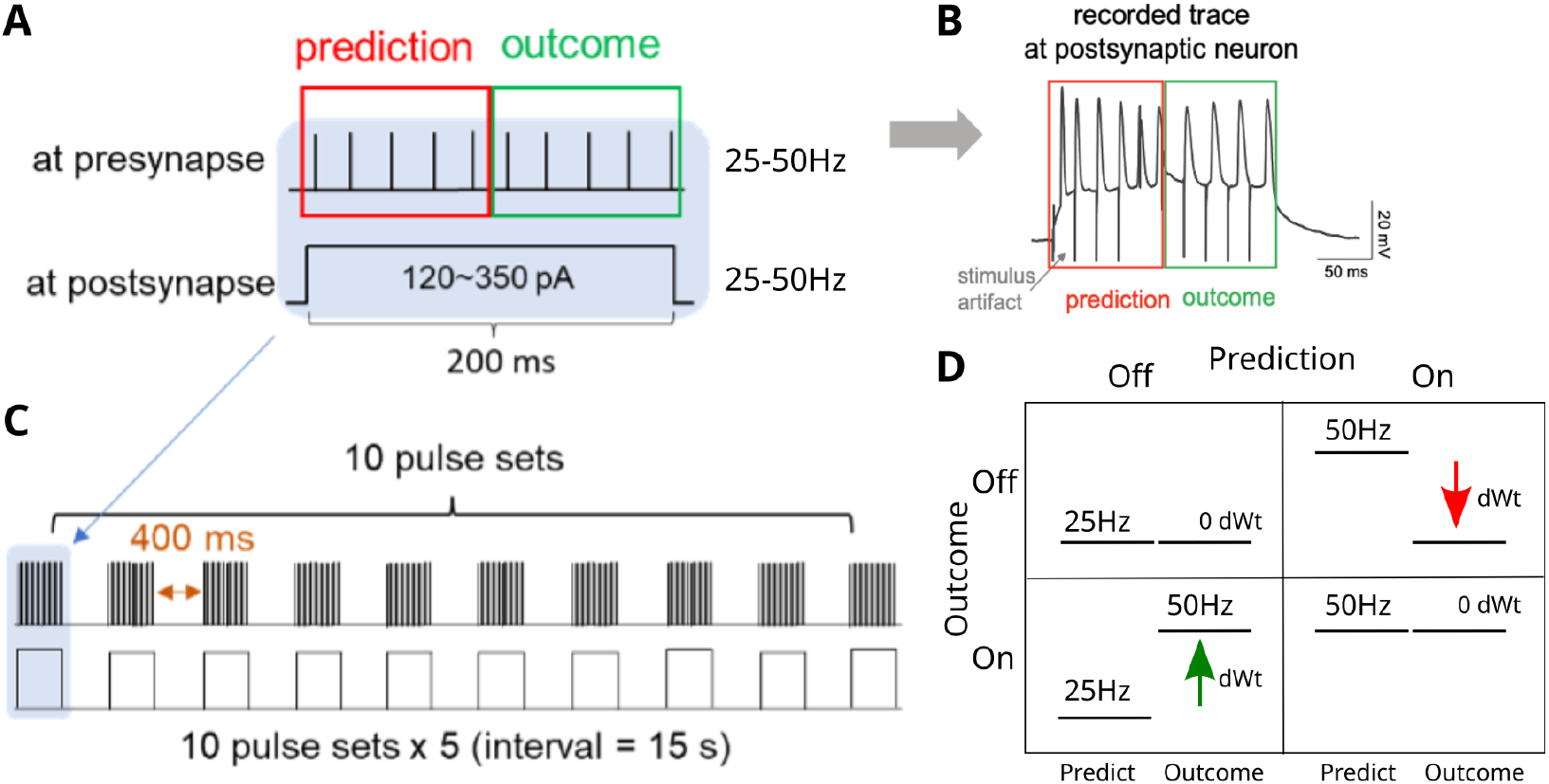
Stimulation protocol to simulate temporal dynamics over a 200 ms theta cycle, where the first 100 ms represents the prediction, and the second 100 ms represents the outcome. **A** Presynaptic activity was driven by direct electrical stimulation of axonal fibers at 25 or 50 Hz, while postsynaptic activity was driven by current clamping at a level that produced the approximate corresponding level of postsynaptic spike rate during initial calibration for each slice (e.g., 120 and 350 pA). **B** Sample trace of postsynaptic membrane potential recorded under patch clamp, under the 50 Hz stable stimulation conditions. **C** The theta-cycle dynamics were repeated 10x with 400 ms spacing, to drive a larger overall synaptic plasticity effect. **D** The predictions from the temporal-derivative learning rule are that LTP (positive delta-weight or dWt) should occur for positive derivatives (outcome – prediction > 0), and LTD (negative dWt) should occur for negative derivatives, while stable activity profiles should result in no net dWt. Note that the stable 50 Hz case has the highest sustained activity level, and yet we predict no weight change, while the two opposite-sign cases have the same overall activity level, and yet we predict different dWt directions. These predictions are inconsistent with standard Hebbian-like mechanisms based on total accumulated activity.

As shown in the figure, there is a 2×2 matrix of prediction (first half) vs. outcome (second half) activity levels, with all combinations of the presynaptic 25 Hz (low) and 50 Hz (high) activity levels coordinated with postsynaptic low and high depolarizations. The temporal-derivative algorithm predicts that a positive temporal change (i.e., outcome – prediction > 0) should result in LTP (long-term potentiation or a positive delta-weight (dWt) change), while a negative temporal change should result in LTD (negative dWt). Furthermore, any stable pattern of activity across the theta cycle should result in no net weight change (0 dWt). This is summarized in the following equation:

**Eq 1:** Temporal derivative learning rule

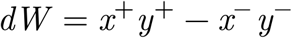

where *x*^+^ is the activity of the sending neuron in the outcome (plus) phase, while *y*^-^ is the activity of the receiving neuron in the prediction (minus) phase, and so forth. This equation can be derived from multiple different starting assumptions (Ackley et al., 1985; Movellan & McClelland, 1993; O’Reilly, 1996), and has been labeled the *Contrastive Hebbian Learning* (*CHL*) equation, because it is the difference or contrast between two Hebbian *xy* factors.

The *contrastive* or temporal derivative aspect of this learning rule is what separates it qualitatively from standard Hebbian learning mechanisms, which generally predict that the direction and magnitude of synaptic plasticity is a function of the overall synaptic activity level, *xy*. Historically, the properties of the NMDA receptor in being sensitive to both pre and postsynaptic activity were quickly recognized to be consistent with the earlier theoretical ideas from Hebb (1949) (Dunwiddie & Lynch, 1978; Lisman, 1989; Bear & Malenka, 1994), with the intracellular calcium levels in the postsynaptic terminal bouton reflecting this Hebbian synaptic activity coproduct. For example, note that the greatest level of overall synaptic activity is in the 50 Hz stable case, where the temporal derivative mechanism predicts 0 weight change, but a standard Hebbian model would predict the greatest level of positive LTP.

As shown in the results below, we found that the direction of synaptic plasticity under our stimulation protocol was entirely consistent with the predictions from the temporal derivative learning rule, and thus strongly inconsistent with a standard Hebbian learning mechanism. We discuss below how a relatively simple competitive binding dynamic between two chemical pathways that have opposite effects on the sign of synaptic changes can produce the temporal derivative learning property. The essential property is that the potentiation pathway has an overall faster response time constant relative to the depression pathway, which then naturally produces a temporal derivative.

## Materials and methods

### Hippocampal slice preparation

Acute hippocampal slices 320 μ*m* thickness were prepared from postnatal day 16-18 C57BL/6 mice using a vibratome (VT 1000S, Leica Microsystems) in ice-cold artificial cerebrospinal fluid (ACSF, in mM: 127 NaCl, 2.5 KCl, 25 Glucose, 25 NaHCO_3_, 1.2 NaH_2_P0_4_, 1 MgCl_2_, 2 CaCl_2_, pH 7.3, oxygenated with 95% O_2_/5% CO_2_). Slices were recovered in ACSF at 32°C for 25-30 min, and then at room temperature for 20-30 min. Slices were used for up to 4-5 hours after recovery.

### Electrophysiology

Electrophysiology was performed using a Multiclamp 700B amplifier (Molecular Devices). Area CA3 of the hippocampus was removed from slices prior to patching. EPSPs were recorded from hippocampal CA1 neurons (P16-18) in current-clamp mode at a sampling rate of 10 kHz filtered at 1 kHz under the whole-cell patch-clamp configurations, in ACSF. For whole-cell patch-clamp recordings, the patch pipettes were filled with (in mM): 136 K-gluconate, 5 NaCl, 10 HEPES, 0.6 EGTA, 4 Na-ATP, 0.4 Na-GTP at pH 7.3 adjusted with KOH. Current injections were performed prior to implementation of the experimental stimulus protocol (Figure 3) to determine the postsynaptic current injection amplitudes for the two phases of the induction protocol, to approximately match the presynaptic 25 and 50 Hz firing rates (e.g., 120 and 350 pA). For the presynaptic stimulation, bipolar platinum-iridium microelectrodes (FHC) were used with an ISO-Flex stimulus isolator. The bipolar electrode was placed in the stratum radiatum approximately 5 mm away from the dendrites of the target CA1 cell. Stimulus strength was adjusted to evoke a ~5 mV EPSP during the 5-6 min baseline recording period. Pipette seal and cell health were monitored throughout the experiment by injection of a short hyperpolarizing current. EPSPs were recorded for up to 45 min after induction. Only one induction was performed per slice.

### Stimulation protocol

To set the stimulation patterns used in our experiments, we first compared the maximum spike rate that could be produced by a stable 200 ms current injection, which was ~55 Hz, with that obtained from driving discrete action potentials using short duration pulses (3 ms), which was ~100 Hz (Figure 4a and b). We chose to move forward with the stable long-duration (200 ms) current injection because that should be more naturalistic. We then determined the length of the gap that was needed between 200 ms theta windows for no obvious degradation across 10 repetitions. As shown in Figure 4c, a gap of 200 ms resulted in degradation of spikes, while a gap of 400 ms did not, so we selected the 400 ms gap.

**Figure 4.**
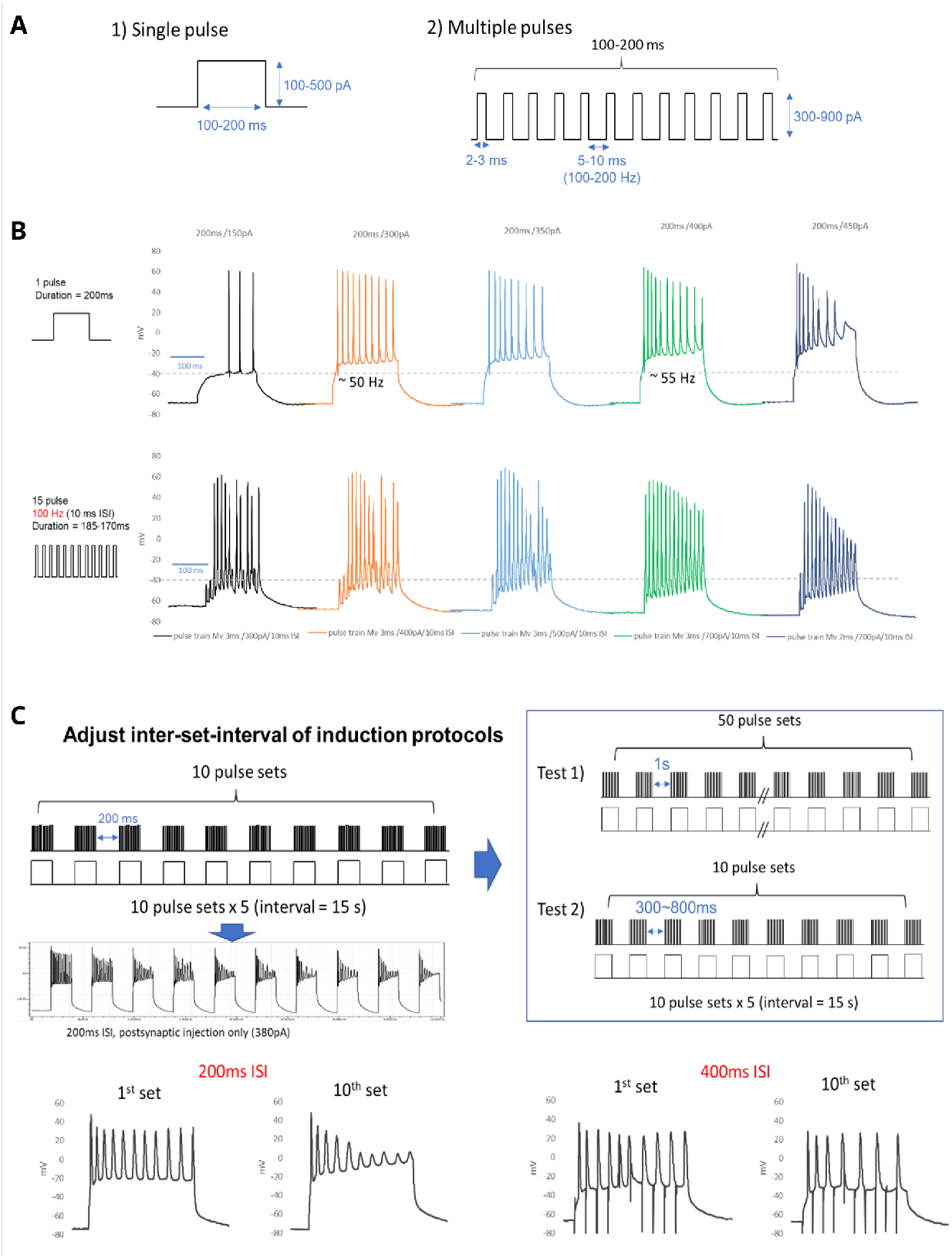
Determination of stimulus parameters. **A** and **B** compare the spiking rates generated by a prolonged postsynaptic current injection across the 200 ms theta cycle, versus discrete short (3 ms) current pulses. The maximum spiking rate from a consistent current was ~55 Hz, with higher currents producing spike dropout, as shown in the upper right plot of panel **B.** Discrete current pulses produced up to 100 Hz firing, but are less representative of the slower postsynaptic depolarizations typically observed in the brain. **C** shows that a 200 ms quiet gap between the 200 ms theta window of patterned activity resulted in visible degradation of spike profile, while 400 ms and above showed no such degradation.

It is important to note that, even after assessing and selecting the appropriate current injection amplitudes to obtain the targeted specific postsynaptic firing rates, we found that postsynaptic spiking rates were often below what was originally expected with a given current injection. We considered it likely that the synapses on the postsynaptic cell respond more to the overall depolarization amplitude and duration for plasticity induction (often in the case of postsynaptic plateau without dendritic spikes altogether), therefore we used fixed low and high current injection amplitudes in our stimulation protocol.

### Data analysis

EPSP amplitude (mV) was measured as the peak membrane depolarization from baseline membrane potential (mean of first 100 ms of recording). Analysis was performed in Clampfit 10.7 (Molecular Devices).

### Statistics

All presented numeric values and graphic representations represent mean ± standard error of the mean, and all statistical analyses used two-tailed t-tests. Statistical significance level (α) was set to p < 0.05 for all tests. A minimum of three independent hippocampal preparations contributed to each data set. All statistics were calculated across cells.

## Results

To test whether synaptic plasticity could be sensitive to the temporal derivative over a period of roughly 200 ms, we manipulated the activity level of CA3 synaptic inputs onto a postsynaptic CA1 neuron, across the two 100 ms halves of the 200 ms theta cycle. Figure 5 shows the results from all four temporal derivative conditions shown in Figure 3, plotting the EPSP amplitude before and after the induction stimulation protocol at time 0. The *both increase* condition (presynaptic neurons stimulated 100 ms at 25 Hz then increased to 50 Hz for the remaining 100 ms, and corresponding postsynaptic low to high depolarization) resulted in LTP (normalized EPSP at 30-35 min = 1.42 ± 0.15, n=12, orange circles and bars in the figure). Conversely, the *both decrease* condition (50 to 25Hz with corresponding high to low depolarization) resulted in LTD (normalized EPSP at 30-35 min = 0.73 ± 0.10, n=18, blue circles and bars in the figure). Finally, both of the flat protocols (constant 25 Hz or 50 Hz with corresponding stable low or high depolarization) resulted in no net change in EPSP amplitudes, and there were no differences between 25 Hz (normalized EPSP at 30-35 min = 1.12 ± 0.15, n=12, light gray circles and bars) and 50 Hz (normalized EPSP at 30-35 min = 1.09 ± 0.08, n=11, dark gray circles and bars).

**Figure 5.**
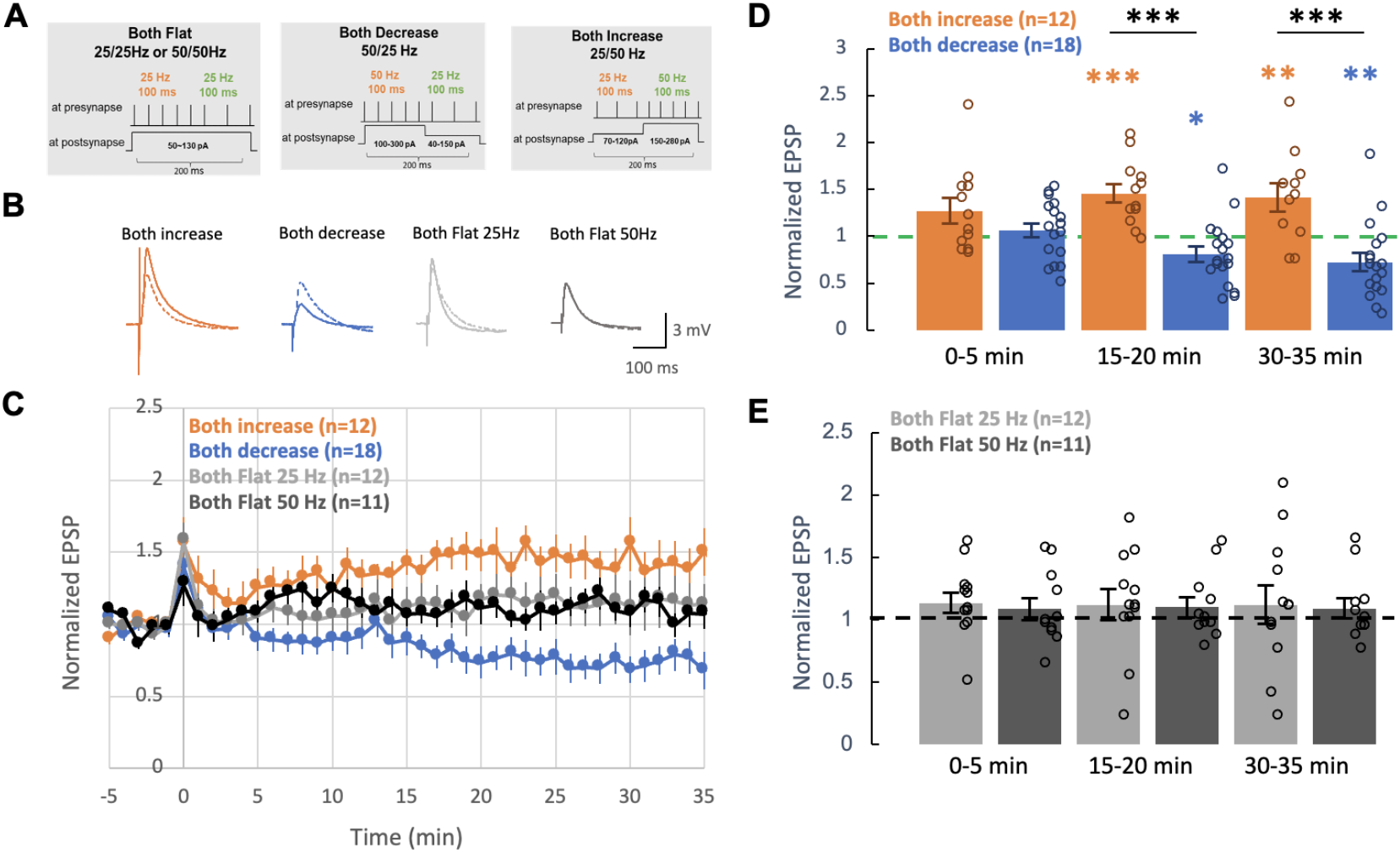
Results, which are consistent with the predictions of the temporal derivative learning mechanism. **A** Schematic of stimulation protocols. **B** Representative traces showing before (dotted line) and after (solid line) for each stimulation protocol. **C** Normalized EPSP amplitudes before and after the stimulation protocol at time 0, showing that the increasing temporal derivative (25 to 50 Hz, low to high, in orange) resulted in LTP, while the decreasing temporal derivative (50 to 25 Hz, high to low, in blue) resulted in LTD. Both flat profiles (constant 25 Hz, low or 50 Hz, high) resulted in no net synaptic efficacy change. **D, E** Summary showing all cell values (open circles) and averages (bars) for the individual conditions at different times after stimulation. n values represent number of cells, with statistically significant results highlighted with asterisks (* = P < .05, ** = P < .01, *** = P < .001).

Our results are fully consistent with the predictions of the temporal derivative learning mechanism (Eq 1), and strongly *inconsistent* with existing Hebbian-like learning rules, which would have predicted that the constant 50 Hz case should exhibit the most LTP, while constant 25 Hz should be the weakest or result in LTD. The idea that the same overall level of spiking, distributed differently across the 200 ms theta window in the 25-to-50 and 50-to-25 cases, could produce opposite patterns of LTP and LTD (respectively) is beyond the scope of standard Hebbian frameworks.

## Discussion

We tested the hypothesis that the direction of synaptic plasticity should be a function of the change in synaptic activity over time, the *temporal derivative*, which would be consistent with the ability to perform an approximation to the computationally powerful error backpropagation learning algorithm. We drove coordinated changes in pre and postsynaptic activity for all four combinations of low *x* high activity (presynaptic firing rates of 25 Hz or 50 Hz, with corresponding low or high postsynaptic depolarization) across two sequential 100 ms windows, and measured the resulting changes in synaptic efficacy on test probes. We found that an increasing change in activity (low to high) resulted in LTP (increased synaptic efficacy), while a decreasing change (high to low) resulted in LTD (decreased synaptic efficacy). Meanwhile, both flat activity profiles, at either a stable low or high level, resulted in no net synaptic efficacy changes.

This pattern of synaptic efficacy changes is entirely consistent with the predictions of the temporal derivative-based approximation to error backpropagation known as *GeneRec* (O’Reilly, 1996; see also Xie & Seung, 2003; Scellier & Bengio, 2017), which is a generalization of the *Recirculation* algorithm (Hinton & McClelland, 1988), and builds on the phase-based learning ideas initially developed in the *Boltzmann Machine* (Ackley et al., 1985). Thus, the results reported here provide critical empirical support for the hypothesis that the neocortex learns via error backpropagation, leveraging the well-established bidirectional excitatory connectivity that is uniquely present in this brain area (Van Essen & Maunsell, 1983; Markov et al., 2013). This bidirectional connectivity allows activity changes in any part of the neocortex to propagate widely, thereby accomplishing the same effect as error backpropagation.

Furthermore, the unique properties of the thalamocortical connectivity between the neocortex and the higher-order thalamic nuclei (pulvinar and mediodorsal) provide a mechanism for reliably driving alternating phases of prediction and outcome activity states (O’Reilly et al., 2021), which is an essential requirement for temporal-derivative based error backpropagation learning. Thus, the available neurobiological data at multiple levels of analysis is overall consistent with the requirements for this computationally powerful form of learning.

### Comparison with Hebbian learning

The predominant computational-level interpretation of neocortical learning in the literature has generally focused on various forms of Hebbian learning, based on the well-established data demonstrating a relationship between the level of postsynaptic calcium, entering via NMDA receptors, and the direction and magnitude of synaptic plasticity (Lisman, 1989; Bear & Malenka, 1994). Specifically, low levels of calcium result in LTD, while higher levels result in LTP. This is generally consistent with the BCM (Bienenstock et al., 1982) version of a Hebbian learning algorithm.

Spike-timing dependent plasticity (STDP) (Bi & Poo, 1998) has been a primary focus of computational models (e.g., Kheradpisheh et al., 2018; Diehl & Cook, 2015). However, it is now clear that the simple computationally compelling form of STDP originally described, which required a very particular stimulation protocol with individual pairs of spikes separated by 1 s intervals, is not generally applicable to more realistic patterns of neural activity (Debanne & Inglebert, 2023). Indeed the same BCM-like pattern emerges with more realistic, denser activity patterns (Shouval et al., 2010; Izhikevich & Desai, 2003).

The critical computational advantage of error backpropagation over these Hebbian learning mechanisms is that it is mathematically designed to coordinate the learning across all of the neurons in the network, to minimize a distal error signal. By contrast, Hebbian learning only has a local, heuristic function in terms of extracting statistical regularities of co-activation (Oja, 1982; Rumelhart & Zipser, 1985; Intrator & Cooper, 1992). Therefore, there is no reason to believe that Hebbian learning can effectively train deep layered networks like those present in the neocortex, whereas this is precisely the case where error backpropagation excels. Thus, the present results provide an important potential way forward in reconciling the computational-level demands of neocortical learning with the underlying neural mechanisms.

More recently, *behavioral timescale synaptic plasticity (BTSP)* has been extensively studied in area CA1 of the hippocampus, where elevated *plateau potentials* in distal dendrites provide the critical plasticity-inducing mechanism that establishes an eligibility trace over the course of several seconds (Magee, 2026; Bittner et al., 2015; Bittner et al., 2017). These distal plateau potentials are activated by entorhinal cortex (EC) layer 3 inputs, which thus serve as a special training signal for driving plasticity in the other major population of synaptic inputs, from area CA3. These EC inputs can encode reward-predictive cues, accounting for the over-representation of rewarded locations in CA1 neurons (Grienberger & Magee, 2022). A similar mechanism has recently been described in layer 5 pyramidal neurons in neocortex, which also have a prominent distal dendritic tuft (Yaeger et al., 2025; Xiao et al., 2025).

In both of these BTSP cases, the learning mechanism appears to be a relatively simple case of rapid and transient plasticity specifically for the *output* pathways, to decode the slowly learning internal representations onto output targets of current behavioral relevance. This is computationally related to the way that *reservoir computing* networks are trained (Verstraeten et al., 2007; Tanaka et al., 2019), where a complex internal dynamical state can be read out by only adapting a single layer of output neuron synapses. There is an inhibitory feedback mechanism that can prevent overlearning in the CA1 neurons (Campbell et al., 2026).

This rapid and transient readout mechanism thus provides a way to preserve slow, incremental learning of systematic internal representations (driven by a version of the computationally powerful error-backpropagation algorithm), while also rapidly adapting to the current behavioral demands, providing a reasonable compromise solution to the fundamental *stability-plasticity dilemma* (Magee, 2026). Specifically, we propose that CA1 neurons, and layer 5 neurons in the neocortex, also exhibit a slower, incremental learning mode as we demonstrate in this paper, along with a rapid BTSP learning mode to specifically focus on areas of greatest behavioral relevance.

### Temporal derivative plasticity via competing kinases

How could the results we obtained arise from the biochemical processes operating at the synapse? Mathematically, the temporal derivative can be computed as the difference between *fast* minus *slow* integrals of a common driving input signal. Intuitively, the fast integral more closely reflects the more recent outcome state, while the slow integral still retains more of the trace from the earlier prediction state. See temporal derivative on compcogneuro.org for an interactive demonstration of this principle.

Although the possible molecular basis for our observations is yet to be determined, we describe one molecular signaling scenario that is well-aligned with the fast-minus-slow mechanism. Neurochemically, existing evidence shows that the difference between LTP versus LTD is determined in part by a competition between two different *kinases, CaMKII* (calcium calmodulin-dependent protein kinase II) and *DAPK1* (death-associated protein kinase 1), both of which are driven by calcium-activated calmodulin (CaM) ((Goodell et al., 2017; Goodell et al., 2021; Cook et al., 2021; Tullis & Bayer, 2023; Bayer & Giese, 2025). If CaMKII had a faster overall integration of the common CaM driver, and DAPK1 a slower such integration, then this would implement the necessary temporal derivative mechanism.

Thus, a difference in overall integration rate stands as a prediction from this overall framework, and is expected to be reflected in the neurochemical processes underlying the form of synaptic plasticity that we report here. There are many questions that remain to be explored, including how the coordination and timing between pre- and postsynaptic activity impacts the outcome.

### Cortical dynamics versus predictive coding

The error-driven learning supported by the corticothalamic prediction vs. outcome mechanism (Figure 2; O’Reilly et al., 2021) represents an alternative to the widely discussed Bayesian predictive coding framework (e.g., Rao & Ballard, 1999; Friston, 2009), and to other proposed implementations of error-backpropagation (Lillicrap et al., 2020). The temporal derivative basis for this alternative greatly simplifies the cortical dynamics necessary to support predictive learning, which aligns better with the available data.

The Bayesian model requires that a sub-population of neurons explicitly represent the *prediction error*, by subtracting a top-down prediction from the bottom-up actual outcome. Thus, different populations of neurons must be somehow segregated so that they can represent fundamentally distinct information. Furthermore, all three of these different signals (prediction, outcome, error) should in principle be communicated across layers, in different directions, requiring strongly segregated pathways.

By contrast, in the temporal derivative model, the entire network is always *coherent* and *synergistic* at any given point in time: all layers and neurons are fundamentally cooperating to represent a consistent interpretation of the *current* state of the world. This current state just alternates over time between representing the prediction versus the outcome. If the outcome matches the prediction, then there is no change, which would typically be the situation in a mature, well-trained system: a stable and accurate representation of the world. However, earlier in developmental learning, and in relatively novel or challenging situations in the mature system, unexpected outcomes can drive learning to improve the accuracy of the prediction states.

This form of learning thus allows for all levels in the network to work together to drive parallel *constraint satisfaction* processing, integrating top-down and bottom-up constraints, to drive coherent interpretations of the current state (Hopfield & Tank, 1985; O’Reilly et al., 2013). This represents a powerful form of search through representation space, operating as a kind of inner-loop optimization within the outer-loop of error backpropagation search through synaptic weight space to improve the predictive accuracy of the system. Computational models reported in O’Reilly et al. (2021) and extensively on compcogneuro.org demonstrate the efficacy of this form of learning and processing, using biologically realistic spiking neurons.

The available neural evidence is consistent with the coherent, synergistic, redundant encoding of information across all levels of the cortex, with no significant evidence of the kind of structural segregation required by the Bayesian model (Walsh et al., 2020; Heilbron & Chait, 2018). The primary positive evidence that has been found, a suppression of neural activity for expected outcomes relative to unexpected ones, is compatible with the alternative temporal derivative model in conjunction with well-established neural adaptation / accommodation mechanisms (Kok & Lange, 2015; see O’Reilly et al., 2021 for detailed discussion).

Thus, the temporal derivative framework supports the widely accepted idea that the neocortex learns by generating top-down predictions of what will happen next, in a way that appears to be more compatible with available neural evidence at multiple levels of analysis.

## Conclusion

In conclusion, the results presented here represent an important first step in testing the possibility that neocortical learning implements the computationally powerful error backpropagation learning mechanism, based on synaptic plasticity that is driven by a temporal derivative. This form of learning is consistent with a wide range of existing data, and would benefit from further experimental investigation that thoroughly tests this possible answer to one of the most important outstanding questions in neuroscience.

## Acknowledgments

This work was funded by the Astera Institute and by the National Institutes of Health (R01 NS137635).

## References

Ackley, D.H., Hinton, G.E., & Sejnowski, T.J. (1985). A learning algorithm for Boltzmann machines. Cognitive Science, 9, 147–169.

Bayer, K.U., & Giese, K.P. (2025). A revised view of the role of CaMKII in learning and memory.Nature Neuroscience, 28, 24–34. https://www.nature.com/articles/s41593-024-01809-x 10.1038/s41593-024-01809-x

Bear, M.F., & Malenka, R.C. (1994). Synaptic plasticity: LTP and LTD. Current Opinion in Neurobiology, 4, 389–399. https://www.sciencedirect.com/science/article/pii/0959438894901015 10.1016/0959-4388(94)90101-5

Bienenstock, E.L., Cooper, L.N., & Munro, P.W. (1982). Theory for the development of neuron selectivity: Orientation specificity and binocular interaction in visual cortex. The Journal of Neuroscience, 2, 32–48. http://www.ncbi.nlm.nih.gov/pubmed/7054394

Bi, G., & Poo, M. (1998). Synaptic modifications in cultured hippocampal neurons: dependence on spike timing, synaptic strength, and postsynaptic cell type. The Journal of Neuroscience, 18, 10464–10472. http://www.jneurosci.org/content/18/24/10464

Bittner, K.C., Grienberger, C., Vaidya, S.P., Milstein, A.D., Macklin, J.J., Suh, J., Tonegawa, S., & Magee, J.C. (2015). Conjunctive input processing drives feature selectivity in hippocampal CA1 neurons. Nature Neuroscience, 18(8), 1133–1142. https://www.nature.com/articles/nn.4062 10.1038/nn.4062

Bittner, K.C., Milstein, A.D., Grienberger, C., Romani, S., & Magee, J.C. (2017). Behavioral time scale synaptic plasticity underlies CA1 place fields. Science, 357, 1033–1036. http://science.sciencemag.org/content/357/6355/1033 10.1126/science.aan3846

Campbell, E.P., Martin, L., Magee, J.C., & Grienberger, C. (2026). Learning-dependent feedback by OLM interneurons shapes CA1 representations. 2025.12.21.695825. https://www.biorxiv.org/content/10.64898/2025.12.21.695825v2 10.64898/2025.12.21.695825

Cook, S.G., Buonarati, O.R., Coultrap, S.J., & Bayer, K.U. (2021). CaMKII holoenzyme mechanisms that govern the LTP versus LTD decision.Science Advances, https://www.science.org/doi/abs/10.1126/sciadv.abe2300 10.1126/sciadv.abe2300

Debanne, D., & Inglebert, Y. (2023). Spike timing-dependent plasticity and memory.Current Opinion in Neurobiology, 80, 102707. https://www.sciencedirect.com/science/article/pii/S0959438823000326 10.1016/j.conb.2023.102707

Diehl, P.U., & Cook, M. (2015). Unsupervised learning of digit recognition using spike-timing-dependent plasticity. Frontiers in Computational Neuroscience, 9, https://www.frontiersin.org/articles/10.3389/fncom.2015.00099/full 10.3389/fncom.2015.00099

Dunwiddie, T., & Lynch, G. (1978). Long-term potentiation and depression of synaptic responses in the rat hippocampus: localization and frequency dependency. The Journal of Physiology, 276, 353–367. https://onlinelibrary.wiley.com/doi/abs/10.1113/jphysiol.1978.sp012239 10.1113/jphysiol.1978.sp012239

Fiebelkorn, I.C., & Kastner, S. (2021). Spike timing in the attention network predicts behavioral outcome prior to target selection. Neuron, 109, 177–188.e4. https://www.sciencedirect.com/science/article/pii/S0896627320307637 10.1016/j.neuron.2020.09.039

Friston, K. (2009). The free-energy principle: a rough guide to the brain? Trends in cognitive sciences, 13, 293–301. http://www.ncbi.nlm.nih.gov/pubmed/19559644

Goodell, D.J., Tullis, J.E., & Bayer, K.U. (2021). Young DAPK1 knockout mice have altered presynaptic function. Journal of Neurophysiology, 125, 1973–1981. http://journals.physiology.org/doi/full/10.1152/jn.00055.2021 10.1152/jn.00055.2021

Goodell, D.J., Zaegel, V., Coultrap, S.J., Hell, J.W., & Bayer, K.U. (2017). DAPK1 mediates LTD by making CaMKII/GluN2B binding LTP specific. Cell Reports, 19, 2231–2243. http://www.sciencedirect.com/science/article/pii/S2211124717307258 10.1016/j.celrep.2017.05.068

Grienberger, C., & Magee, J.C. (2022). Entorhinal cortex directs learning-related changes in CA1 representations. Nature, 611, 554–562. https://www.nature.com/articles/s41586-022-05378-6 10.1038/s41586-022-05378-6

Hebb, D.O. (1949). The Organization of Behavior. Wiley.

Heilbron, M., & Chait, M. (2018). Great Expectations: Is there Evidence for Predictive Coding in Auditory Cortex? Neuroscience, 389, 54–73. https://www.sciencedirect.com/science/article/pii/S030645221730547X 10.1016/j.neuroscience.2017.07.061

Hinton, G.E., & McClelland, J.L. (1988). Learning representations by recirculation. In D.Z. Anderson (Ed.), Neural Information Processing Systems (NIPS 1987 (pp. 358–366)) American Institute of Physics. http://papers.nips.cc/paper/78-learning-representations-by-recirculation.pdf

Hopfield, J.J., & Tank, D.W. (1985). {`Neural’} computation of decisions in optimization problems. Biological Cybernetics, 52, 141–152. http://www.ncbi.nlm.nih.gov/pubmed/4027280

Intrator, N., & Cooper, L.N. (1992). Objective function formulation of the BCM theory of visual cortical plasticity: Statistical connections, stability conditions. Neural Networks, 5, 3–17. https://www.sciencedirect.com/science/article/pii/S0893608005800036 10.1016/S0893-6080(05)80003-6

Izhikevich, E.M., & Desai, N.S. (2003). Relating STDP to BCM. Neural computation, 15, 1511–1524. http://www.ncbi.nlm.nih.gov/pubmed/12816564

Ketz, N., Morkonda, S.G., & O’Reilly, R.C. (2013). Theta coordinated error-driven learning in the hippocampus. PLoS Computational Biology, 9, e1003067. http://www.ncbi.nlm.nih.gov/pubmed/23762019

Kheradpisheh, S.R., Ganjtabesh, M., Thorpe, S.J., & Masquelier, T. (2018). STDP-based spiking deep convolutional neural networks for object recognition. Neural Networks, 99, 56–67. https://www.sciencedirect.com/science/article/pii/S0893608017302903 10.1016/j.neunet.2017.12.005

Kok, P., & Lange, F.P. (2015).Predictive Coding in Sensory Cortex. In An Introduction to Model-Based Cognitive Neuroscience (pp. 221–244). Springer, New York, NY. https://link.springer.com/chapter/10.1007/978-1-4939-2236-9_11 10.1007/978-1-4939-2236-9_11

Lillicrap, T.P., Santoro, A., Marris, L., Akerman, C.J., & Hinton, G. (2020). Backpropagation and the brain. Nature Reviews Neuroscience, 21(6), 335–346. https://www.nature.com/articles/s41583-020-0277-3 10.1038/s41583-020-0277-3

Lisman, J. (1989). A mechanism for the Hebb and the anti-Hebb processes underlying learning and memory. Proceedings of the National Academy of Sciences, 86, 9574–9578. https://www.pnas.org/doi/abs/10.1073/pnas.86.23.9574 10.1073/pnas.86.23.9574

Magee, J.C. (2026). Behavioral timescale synaptic plasticity: properties, elements and functions. Nature Neuroscience, 29, 520–534. https://www.nature.com/articles/s41593-026-02214-2 10.1038/s41593-026-02214-2

Markov, N.T., Ercsey-Ravasz, M., Lamy, C., Ribeiro Gomes, A.R., Magrou, L., Misery, P., Giroud, P., Barone, P., Dehay, C., Toroczkai, Z., Knoblauch, K., Van Essen, D.C., & Kennedy, H. (2013). The role of long-range connections on the specificity of the macaque interareal cortical network. Proceedings of the National Academy of Sciences U. S. A., 110, 5187–5192. http://www.ncbi.nlm.nih.gov/pubmed/23479610

Montague, P.R., Dayan, P., & Sejnowski, T.J. (1996). A framework for mesencephalic dopamine systems based on predictive Hebbian learning. Journal of Neuroscience, 16, 1936–1947. http://www.ncbi.nlm.nih.gov/pubmed/8774460

Movellan, J.R., & McClelland, J.L. (1993). Learning Continuous Probability Distributions with Symmetric Diffusion Networks. Cognitive Science, 17, 463–496.

Oja, E. (1982). A simplified neuron model as a principal component analyzer.Journal of mathematical biology, 15, 267–273. http://www.ncbi.nlm.nih.gov/pubmed/7153672

O’Reilly, R.C. (1996). Biologically plausible error-driven learning using local activation differences: The generalized recirculation algorithm. Neural Computation, 8, 895–938. https://www.mitpressjournals.org/doi/abs/10.1162/neco.1996.8.5.895 10.1162/neco.1996.8.5.895

O’Reilly, R.C., Russin, J.L., Zolfaghar, M., & Rohrlich, J. (2021). Deep Predictive Learning in Neocortex and Pulvinar. Journal of Cognitive Neuroscience, 33, 1158–1196. https://doi.org/10.1162/jocn_a_01708 10.1162/jocn_a_01708

O’Reilly, R.C., Wyatte, D., Herd, S.A., Mingus, B., & Jilk, D.J. (2013). Recurrent Processing during Object Recognition. Frontiers in Psychology, 4, http://www.ncbi.nlm.nih.gov/pubmed/23554596

Rao, R.P., & Ballard, D.H. (1999). Predictive coding in the visual cortex: A functional interpretation of some extra-classical receptive-field effects. Nature Neuroscience, 2, 79–87. http://www.ncbi.nlm.nih.gov/pubmed/10195184 10.1038/4580

Rumelhart, D.E., Hinton, G.E., & Williams, R.J. (1986). Learning representations by back-propagating errors. Nature, 323, 533–536.

Rumelhart, D.E., & Zipser, D. (1985). Feature discovery by competitive learning* Cognitive Science, 9, 75–112. http://onlinelibrary.wiley.com/doi/10.1207/s15516709cog0901_5/abstract 10.1207/s15516709cog0901_5

Scellier, B., & Bengio, Y. (2017). Equilibrium propagation: Bridging the gap between energy-based models and backpropagation. Frontiers in Computational Neuroscience, 11, http://www.ncbi.nlm.nih.gov/pmc/articles/PMC5415673/ 10.3389/fncom.2017.00024

Sherman, S.M., & Guillery, R.W. (2006). Exploring the Thalamus and Its Role in Cortical Function. MIT Press. http://www.scholarpedia.org/article/Thalamus

Sherman, S.M., & Usrey, W.M. (2024). Transthalamic Pathways for Cortical Function.Journal of Neuroscience, 44, https://www.jneurosci.org/content/44/35/e0909242024 10.1523/JNEUROSCI.0909-24.2024

Shouval, H.Z., Wang, S.S., & Wittenberg, G.M. (2010). Spike timing dependent plasticity: A consequence of more fundamental learning rules. Frontiers in Computational Neuroscience, 4, http://www.ncbi.nlm.nih.gov/pubmed/20725599

Sutton, R.S., & Barto, A.G. (1998). Reinforcement Learning: An Introduction. MIT Press. http://www.cs.ualberta.ca/sutton/book/ebook/the-book.html

Tanaka, G., Yamane, T., Héroux, J.B., Nakane, R., Kanazawa, N., Takeda, S., Numata, H., Nakano, D., & Hirose, A. (2019). Recent advances in physical reservoir computing: A review. Neural Networks, 115, 100–123. https://www.sciencedirect.com/science/article/pii/S0893608019300784 10.1016/j.neunet.2019.03.005

Tullis, J.E., & Bayer, K.U. (2023). Distinct synaptic pools of DAPK1 differentially regulate activity-dependent synaptic CaMKII accumulation. iScience, 26, https://www.cell.com/iscience/abstract/S2589-0042(23)00800-3 10.1016/j.isci.2023.106723

Van Essen, D.C., & Maunsell, J.H.R. (1983). Hierarchical organization and functional streams in the visual cortex. Trends in Neurosciences, 6, 370–375.

Verstraeten, D., Schrauwen, B., D’Haene, M., & Stroobandt, D. (2007). An experimental unification of reservoir computing methods. Neural Networks, 20, 391–403. http://www.ncbi.nlm.nih.gov/pubmed/17517492 10.1016/j.neunet.2007.04.003

Walsh, K.S., McGovern, D.P., Clark, A., & O’Connell, R.G. (2020). Evaluating the neurophysiological evidence for predictive processing as a model of perception. Annals of the New York Academy of Sciences, 1464, 242–268. https://www.ncbi.nlm.nih.gov/pmc/articles/PMC7187369/ 10.1111/nyas.14321

Werbos, P. (1974). Beyond Regression: New Tools for Prediction and Analysis in the Behavioral Sciences. [unpublished thesis, Harvard University].

Widrow, B., & Hoff, M.E. (1960). Adaptive Switching Circuits. In Institute of Radio Engineers, Western Electronic Show and Convention, Convention Record, Part 4 (pp. 96–104).

Xiao, K., Li, Y., Sullivan, B.J., Li, G., & Magee, J.C. (2025). Rapid neocortical network modifications via dendritic plateau potential induced plasticity. 2025.11.19.689338. https://www.biorxiv.org/content/10.1101/2025.11.19.689338v1 10.1101/2025.11.19.689338

Xie, X., & Seung, H.S. (2003). Equivalence of backpropagation and Contrastive Hebbian Learning in a layered network. Neural Computation, 15, 441–454. http://www.ncbi.nlm.nih.gov/pubmed/12590814

Yaeger, C.E., Soto-Albors, R.M., Liu, W., Beltramini, A., & Harnett, M.T. (2025). Plateau potentials are instructive signals for behavioral timescale synaptic plasticity in the neocortex. 2025.11.07.687250. https://www.biorxiv.org/content/10.1101/2025.11.07.687250v1 10.1101/2025.11.07.687250

Zheng, Y., Liu, X.L., Nishiyama, S., Ranganath, C., & O’Reilly, R.C. (2022). Correcting the hebbian mistake: Toward a fully error-driven hippocampus. PLOS Computational Biology, 18, e1010589. https://journals.plos.org/ploscompbiol/article?id=10.1371/journal.pcbi.1010589 10.1371/journal.pcbi.1010589

